# Confined migration promotes cancer metastasis through resistance to anoikis and increased invasiveness

**DOI:** 10.1101/2021.08.17.456592

**Authors:** Deborah Fanfone, Zhi Chong Wu, Jade Mammi, Kevin Berthenet, David Neves, Kathrin Weber, Andrea Halaburkova, François Virard, Félix Bunel, Hector Hernandez-Vargas, Stephen WG Tait, Anca Hennino, Gabriel Ichim

**Author notes:** Corresponding author: Gabriel Ichim (<u>mailto:</u>).

## Abstract

Mechanical stress is known to fuel several hallmarks of cancer, ranging from genome instability to uncontrolled proliferation or invasion. Cancer cells are constantly challenged by mechanical stresses not only in the primary tumour but also during metastasis. However, this latter has seldom been studied with regards to mechanobiology, in particular resistance to anoikis, a cell death programme triggered by loss of cell adhesion. Here, we show *in vitro* that migrating breast cancer cells develop resistance to anoikis following their passage through microporous membranes mimicking confined migration (CM), a mechanical constriction that cancer cells encounter during metastasis. This CM-induced resistance was mediated by Inhibitory of Apoptosis Proteins (IAPs), and sensitivity to anoikis could be restored after their inhibition using SMAC mimetics. Anoikis-resistant mechanically-stressed cancer cells displayed enhanced cell motility and evasion from natural killer cell-mediated immune surveillance, as well as a marked advantage to form lung metastatic lesions in mice. Our findings reveal that CM increases the metastatic potential of breast cancer cells.

## Introduction

The majority of cancer-related deaths arise following metastasis ^1 2^. Metastatic cancer cells acquire *de novo* phenotypic traits allowing them to efficiently leave the primary tumour, enter blood circulation and survive harsh conditions, then exit the bloodstream and establish metastasis at a distant site. Though the mechanisms driving cancer cell migration and invasion are well documented, with a clear understanding of epithelial-mesenchymal transition and the metastatic niche ^3^, no efficient therapeutic strategies currently prevent metastasis formation. Metastasis is thus largely incurable, yet this lengthy process can take years and the therapeutic window is therefore large enough to envisage its targeting ^4^.

Multiple mechanical forces occur at each step of cancer development, from the primary tumour to metastasis. During early tumour development, excessive cell proliferation, massive extracellular matrix deposition or cancer-associated fibroblasts exert compressive forces on cancer cells that can reach up to 10 kPa in pancreatic ductal adenocarcinoma ^5^. As they engage in their metastatic journey, cancer cells migrate through barriers including desmoplastic tumour stroma, basal membranes, endothelial layers, and when entering low-diameter capillaries. In healthy tissues or tumours, the extracellular matrix creates pores or tunnel-shaped tracks that are often smaller than the diameter of a cell and migrating cells adjust their shape and size according to these constrictions ^6^. These events, known as confined migration (CM), have dramatic consequences on cancer cells and might even break the nuclear envelope and trigger DNA damage and mutagenesis if the breaks are not efficiently repaired ^7 8^.

CM can alter the phenotypic traits of cancer cells rendering them even more aggressive. More precisely, repeated nuclear deformations and loss of nuclear envelope integrity can activate an invasive programme, and engage the pro-oncogenic Ras/MAPK signalling pathway ^9 10 11^. Once inside the blood or the lymphatic vessels, circulating tumour cells (CTCs) are confronted with deadly fluid shear stress until they extravasate ^12^. Moreover, most CTCs are eliminated by apoptosis in a process called anoikis, occurring when cells detach from the extracellular matrix ^13 14^. By rapidly engaging either the death receptors or the mitochondrial pathway of apoptosis, anoikis has evolved as an efficient physiological barrier for preventing the formation of metastatic colonies by CTCs reaching target organs ^13^. Nonetheless, cancer cells developed strategies to evade anoikis such as overly activated Ras/ERK and PI3K/Akt pathways, engaging the tyrosine kinase receptor TrkB, inactivation of E-cadherin and p53 or enhanced autophagy ^15 16 17 18 19^.

Mechanical stress has emerged as a key factor in shaping the pro-metastatic features of cancer cells. We thus hypothesized that CM may also contribute to the metastatic potential by impacting anoikis and cancer invasiveness. We show here that breast cancer cells having undergone CM, but not compression, become resistant to anoikis, through a mechanism involving lowering apoptotic caspase activation through an upregulation of Inhibitory of Apoptosis Proteins (IAPs). We also report that treatment with SMAC mimetics to lower IAP expression restores the sensitivity to anoikis. Ultimately, a single round of CM is sufficient to enhance emerging aggressiveness, the most obvious effects observed being random migration and escape from natural killer cell-mediated immune surveillance. In addition, these observations are endorsed *in vivo* by higher lung metastatic burden when mice are engrafted with breast cancer cells challenged by CM. Taken together, our results support that CM triggers a particular signalling signature that might favour certain metastatic hallmarks such as resistance to anoikis and increased invasiveness.

## Results

### Confined migration confers breast cancer cells with resistance to anoikis

The human breast cancer cells, MDA-MB-231, are highly invasive and aggressive *in vitro* and *in vivo*, and represent an ideal cellular model to study metastasis ^20^. To investigate the effects of constriction on these cells, we subjected them to a forced passage through a membrane with 3 µm in diameter pores, via a serum gradient, mimicking the confined migration (CM) encountered during cancer progression ^10 21 22^. MDA-MB-231 cells were seeded onto a matrigel-coated tissue culture insert, prior to applying the serum gradient, and thus initially invaded the matrigel plug before following the serum through the microporous membrane (**Fig. 1a**). We ascertained that CM did not affect cell viability, by verifying the incorporation of Calcein AM (viability dye), and by showing that apoptosis-triggering cytochrome *c* was not released by the mitochondria of CM cells, as it co-localized with COX IV (mitochondrial marker) (**Fig. 1b, c**). Since caspases are the main apoptotic executioners, we next tested if they were activated in CM-challenged cancer cells. For this experiment, MDA-MB-231 cells expressing a bimolecular fluorescence complementation (BiFC)-based caspase-3 reporter ^23^, which is functional in actinomycin D-treated and not in CRISPR^BAX/BAK^ cells, were subjected to CM (**Supplementary Fig. 1a, b**). Recovered cells were viable and had not activated apoptotic effector caspases (**Fig. 1d**), substantiating our previous result. In line with this, CM MDA-MB-231 cells displayed an unaltered mitochondrial membrane potential, ATP production and did not generate excessive reactive oxygen species (ROS) (**Supplementary Fig. 1c-f**). Moreover, these cells had a comparable proliferation rate and cell cycle as control cells, as determined by InCucyte-based live-cell microscopy and flow cytometry (**Fig. 1e and Supplementary Fig. 1g**). Hence, the CM model used here did not alter cell viability and was deemed suitable for studying phenotypic changes occurring in these mechanically-challenged cancer cells.

**Figure 1:**
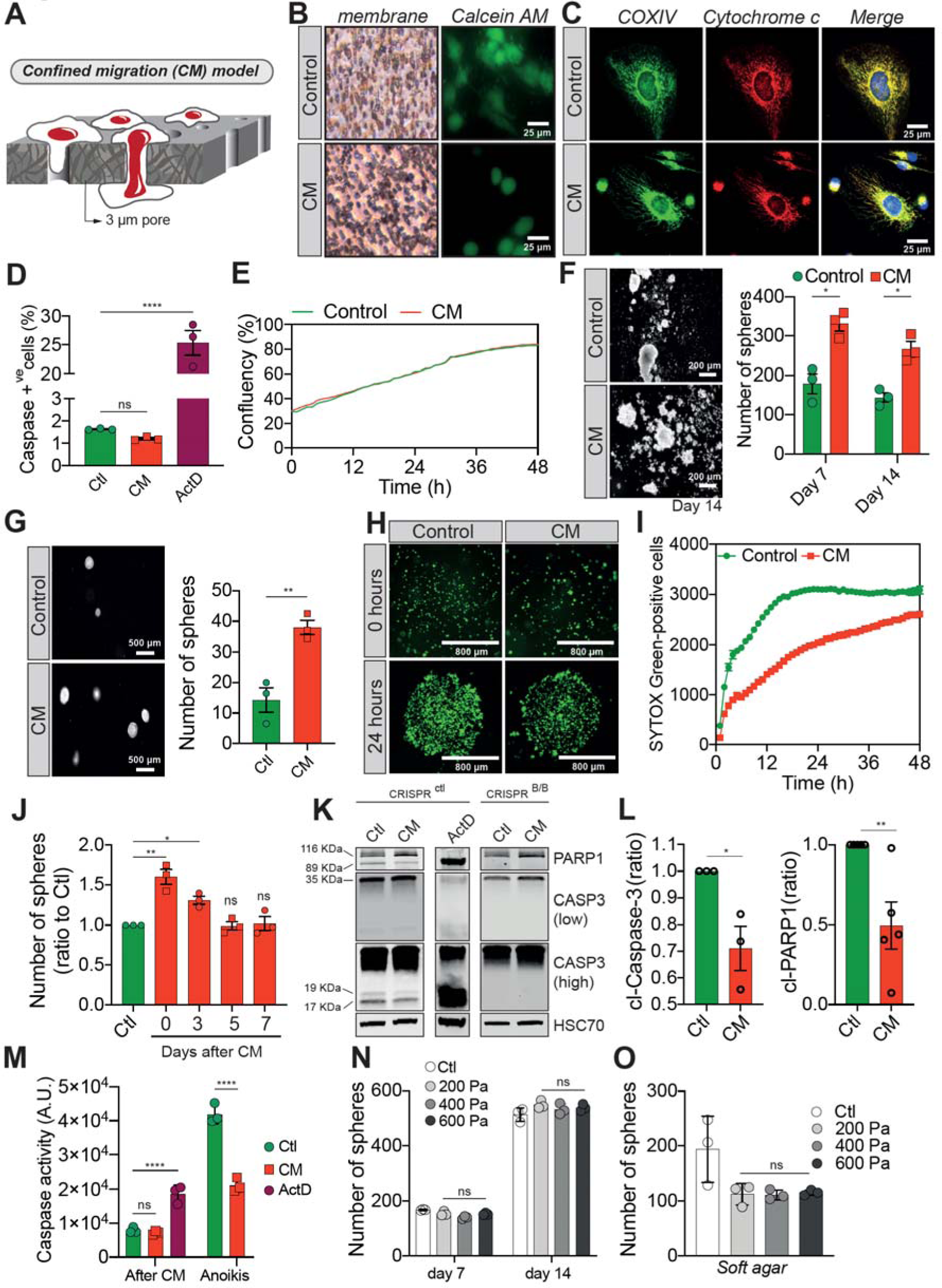
Confined migration confers breast cancer cells with resistance to anoikis. a) Schematic illustration of the 3 µm transwell-based confined migration (CM) model. b) MDA-MB-231 control cells or cells recovered from CM were stained with the Calcein AM viability dye and imaged by epifluorescence microscopy. c) Representative immunofluorescence images of control and CM-challenged MDA-MB-231 cells stained for COX IV and cytochrome *c*. d) Flow cytometry-based quantitative analysis of cells activating the VC3AI caspase reporter (n = 3, One-way ANOVA statistical test). e) IncuCyte ZOOM live-cell imaging-based analysis of cell proliferation (n=3, a representative experiment is shown). f) Representative images of clonogenic structures from control and confined MDA-MB-231 cells grown in anoikis-promoting, ultra-low attachment conditions (left panel). Corresponding quantification of resistance to anoikis after 7 and 14 days of culture (right panel, n = 3, Two- way ANOVA statistical test). g) Control and CM-challenged MDA-MB-231 cells were grown in soft agar to test their anchorage-independent growth. Left panel depicts representative images, while the right panel is the quantification of clonogenic structures after 1 month (n = 3, t test). h) Control and CM-challenged MDA-MB-231 cells were imaged for 48 h in an IncuCyte ZOOM imager in anoikis-promoting, ultra-low attachment conditions, and stained with SYTOX Green (n=3, a representative experiment is shown). i) IncuCyte-based SYTOX Green staining quantification of cell survival in control and CM MDA-MB-231 cells in anoikis-promoting conditions (n=3, a representative experiment is shown). j) Quantitative analysis of clonogenic structures illustrating the duration of resistance to anoikis between control and CM MDA-MB-231 cells up to 7 days post-CM (n = 3, One-way ANOVA statistical test). k) Western blot analysis of PARP-1 cleavage and caspase-3 processing following confined migration and anoikis growth in control and in CRISPR/Cas9-mediated BAX/BAK DKO MDA-MB-231 cells. Actinomycin D treatment (1 µM for 12 h) is used as a positive control for induction of apoptosis. l) Densitometry analysis of PARP-1 and caspase-3 cleavage (ratio of CM cells to control) in anoikis conditions in MDA-MB-231 cells (n = 3-4, t test). m) The effect of CM on effector caspase activation was assessed using a fluorometric assay. Caspase activation was tested either immediately after CM, either in cells that were subsequently grown 24 h in anoikis conditions (n = 3, Two-way ANOVA statistical test). n) Quantification of clonogenic structures formed by control or compressed MDA-MB-231 cells (subjected to 200, 400 or 600 Pa of compression for 16 h) after 7 and 14 days of culture in anoikis conditions (n = 3, Two-way ANOVA statistical test). o) Quantification of clonogenic structures formed in soft agar by compressed MDA-MB-231 cells (n = 3, One-way ANOVA statistical test).

We focused on their response to anoikis, as we hypothesized that circulating tumour cells (CTCs) capable of withstanding such a physiological barrier ^13^ and forming metastases may have acquired tumourigenic properties through the unique mechanical constriction imposed by CM. Consistently, CM-challenged MDA-MB-231 cells survived, grew and formed clonogenic structures in low-attachment conditions, as evidenced by the spheres generated, more efficiently than control cells (**Fig. 1f)**. The effect was not restricted to this cell line since CM-challenged Hs578T breast cancer cells also developed more colonies than control counterparts, indicating that CM-challenged cells had overcome anoikis (**Supplementary Fig. 1h**). This is also the case when cells are grown in soft agar, in an anchorage-independent manner (**Fig. 1g**). In addition, resistance to anoikis was assessed by IncuCyte Imager-based real-time imaging, using SYTOX Green dye exclusion, and again breast cancer cells undergoing CM had a survival advantage when grown in low-attachment conditions (**Fig. 1h, i**). Next, we investigated whether the survival advantage acquired following a single round of CM was transient. MDA-MB-231 cells were challenged by CM once and anoikis resistance was quantified at 3, 5 and 7 days post-CM, revealing that resistance to anoikis was transient (**Fig. 1j**). Since anoikis is a variant of apoptosis, we wondered whether the activation of pro-apoptotic effector caspases was affected by CM. This was assessed by immunoblotting for cleaved caspase-3 and PARP-1, a proxy for efficient caspase activation. Strikingly, CM-challenged cancer cells had lower caspase-3 processing into the active p17 and p19 fragments, whereas PARP-1 cleavage followed the same pattern (**Fig. 1k, l**). Inhibition of effector caspase activation was also confirmed using a fluorometric caspase-3/7 assay, demonstrating that inhibition was particularly important for cells grown in ultra-low attachment and soft agar conditions (further designated as anoikis- favouring conditions) (**Fig. 1m**).

To verify whether this resistance to anoikis was specific to cells having undergone CM and could not arise following the compressive stress experienced within primary tumours upon uncontrolled proliferation or increased extracellular matrix deposition, we tested the effects of compression on resistance to anoikis. We exposed MDA-MB-231 cells *in vitro* to a defined compression by pressing them against a permeable membrane with a weighted piston. The different weights translated into different pressures (200, 400 or 600 Pa) (**Supplementary Fig. 1i**). As shown by the increased nuclear size in compressed cells, this device was suitable to evaluate the effects of compression (**Supplementary Fig. 1j**). Resistance to anoikis, as assessed by growing these cells in anoikis-favouring conditions, was not modified under compression (**Fig. 1n, o**), suggesting that acquisition of resistance to anoikis may be specific to CM. In addition, cancer cell migration through 8 µm in diameter microporous transwells, which do not impose cellular constriction, did not confer cancer cells with resistance to anoikis (**Supplementary Fig. 1k, l**) and had no impact on caspase activation (**Supplementary Fig. 1m**), thus uncoupling cell chemotaxis from this resistance.

In conclusion, these results show that confined migration has a profound impact on cancer cells resistance to cell death through inhibition of pro-apoptotic caspases.

### Confined migration-driven resistance to anoikis relies on the anti-apoptotic IAP proteins

Previous studies reported that Inhibitory of Apoptosis Proteins (IAPs) such as cIAP1, cIAP2 and XIAP promote resistance to anoikis in several cancers, through caspase inhibition ^24 25 26^. We therefore hypothesized that under CM challenge, IAP expression may underlie resistance to anoikis. This was tested by immunoblotting for cIAP1, cIAP2 and XIAP protein expression in CM MDA-MB-231 cells grown in anoikis-favouring conditions, which revealed an upregulation of all three IAPs (**Fig. 2a,** left panel for densitometry analysis). Conversely, MDA-MB-231 cells subjected to compression or migration through 8 µm in diameter microporous transwells had unaltered levels of IAPs (**Supplementary Fig. 2a, b**). To further investigate this correlation, we transiently overexpressed all three IAPs in MDA-MB-231 cells (**Fig. 2b**). Enforced expression of cIAP1 and XIAP significantly enhanced resistance to anoikis in cells grown under anoikis-favouring conditions, while cIAP1 overexpression was dispensable (**Fig. 2c-e**). In a complementary set of experiments, we deleted all three IAPs in MDA-MB-231 cells through CRISPR/Cas9-mediated gene editing (**Fig. 2f**). Following CM and growth in anoikis-favouring conditions, control cells (EV) displayed the expected resistance to anoikis, whereas IAP-depleted cells lost their survival advantage (**Fig. 2g and Supplementary Fig. 2c**). In addition, the use of a SMAC mimetic, one of several developed to specifically induce IAP degradation, namely BV6 ^27^, successfully depleted both cIAP1 and XIAP (**Fig. 2h**). Interestingly, it also abrogated the resistance to anoikis observed in CM- challenged breast cancer cells (**Fig. 2i, j**).

**Figure 2:**
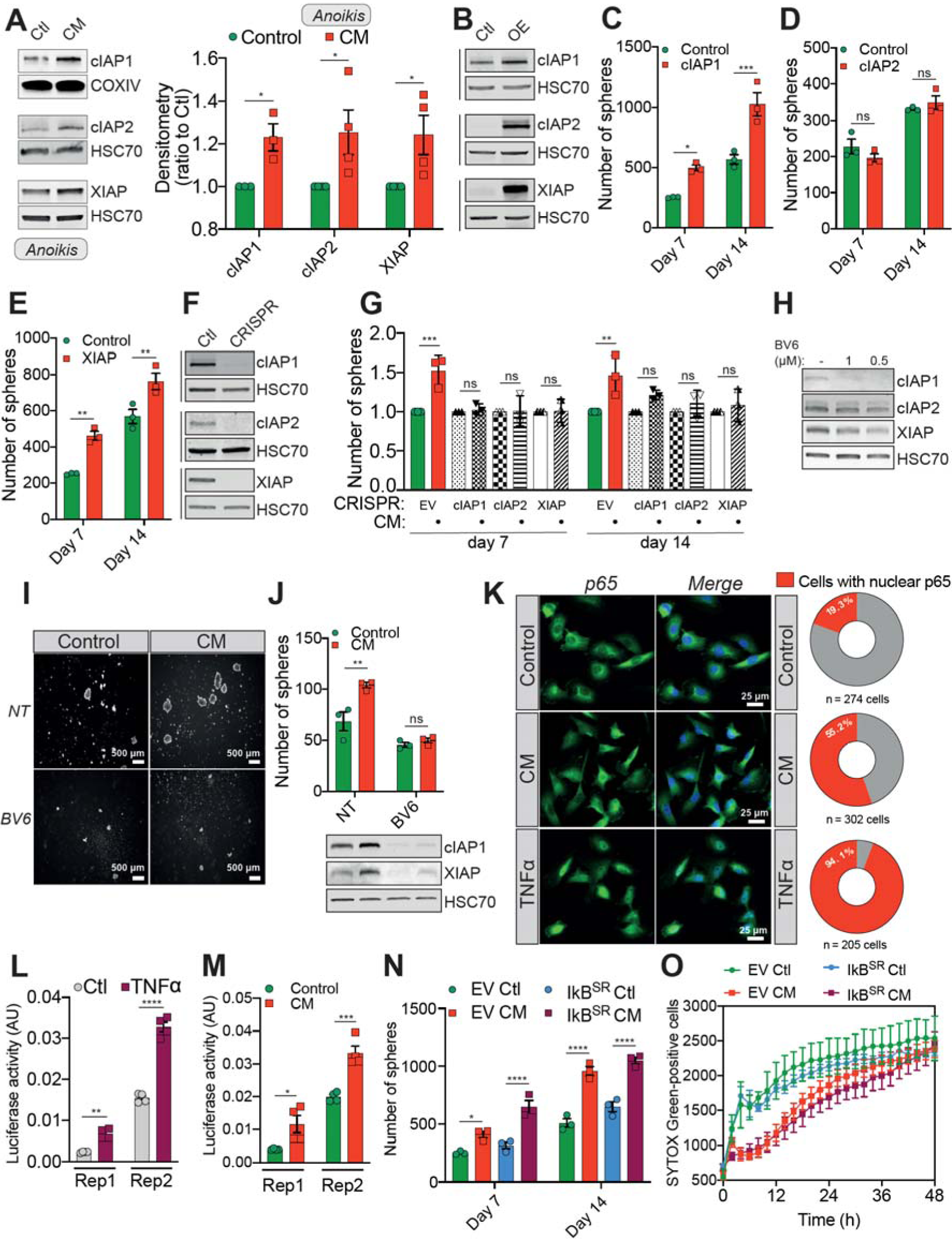
Confined migration-induced resistance to anoikis relies on IAPs. a) Western blot analysis of IAP protein expression in MDA-MB-231 cells after confined migration (CM) through 3 µm in diameter membranes, with cells grown in ultra-low attachment conditions (left panel) and the corresponding densitometry analysis (right panel) (n = 3-4, Two-way ANOVA statistical test). b) Validation of IAP protein overexpression in MDA-MB-231 cells by Western blot. c-e) Quantitative comparison of resistance to anoikis in c-IAP1 (c), cIAP2 (d) and XIAP (e) overexpressing MDA-MB-231 cells in steady-state conditions. Clonogenic structures were counted after 7 and 14 days of culture in ultra-low attachment condition (n = 3, Two-way ANOVA statistical test). f) Western blot analysis of protein expression validating the efficacy of CRISPR/Cas9- mediated deletion of IAP proteins. g) Quantitative comparison of resistance to anoikis in MDA-MB-231 cells deleted for cIAP1, cIAP2 and XIAP using CRISPR/Cas9 and subjected to CM (n = 3, Two-way ANOVA statistical test). h) Validation by Western blot of IAP expression inhibition by treating MDA-MB-231 cells with BV6. Two concentrations of BV6 (0.5 and 1 µM) were used for 16 h of treatment. i) Representative images of clonogenic structures from MDA-MB-231 control and CM cells cultured in low attachment conditions and treated with 0.5 µM of BV6. j) Quantitative comparison of resistance to anoikis in CM MDA-MB-231 cells treated with 0.5 µM of SMAC mimetic BV6. Western blot analysis confirming inhibition of cIAP1 and XIAP expression following BV6 treatment (n = 3, Two-way ANOVA statistical test). k) The effect of confined migration on p65 nuclear localization in MDA-MB-231 cells was assessed by immunofluorescence. TNFα treatment (20 ng/mL for 1 h) was used to induce activation and nuclear translocation of p65. Corresponding quantification of the percentage of cells with nuclear p65 staining is shown in the right panel. l) Luciferase assay to validate the NFkB luciferase reporters following stimulation with TNFα (20 ng/mL for 1 h) (n = 4, Two-way ANOVA statistical test). m) MDA-MB-231 cells transiently expressing two NFkB luciferase reporters were mechanically stressed by CM and luciferase activity was immediately assessed (n = 3, Two- way ANOVA statistical test). n) Empty vector or IkB^SR^-expressing MDA-MB-231 control or CM cells were cultured for 7 and 14 days in anoikis-favouring conditions and the surviving clonogenic structures were quantified (n = 3, Two-way ANOVA statistical test). o) Quantification of resistance to anoikis in empty vector or IkB^SR^-expressing MDA-MB-231 by SYTOX Green incorporation and IncuCyte ZOOM-based live-cell microscopy (n=3, a representative experiment is shown).

Next, we sought to understand the mechanisms responsible for IAP upregulation in constricted cells. We initially performed qRT-PCR to assess IAP mRNA expression in cells subjected to CM, and uncovered that their expression was comparable to control cells, suggesting that IAP expression may be regulated at the post-transcriptional level (**Supplementary Fig. 2d**). IAP contain a RING domain with E3 ubiquitin ligase activities, mediating their own K48 polyubiquitination and that of other protein targets which, is crucial for their role in suppressing apoptosis ^28^. To identify a possible post-transcriptional regulation of IAPs, we first compared the total amount of K48-ubiquitin-linked proteins in control and CM cells. Constricted cells displayed an accumulation of ubiquitinated proteins, indicating a possible bottleneck for protein degradation in CM-stressed cells (**Supplementary Fig. 2e**). In addition, we performed a chase assay with the protein synthesis inhibitor cycloheximide, in order to assess protein half-life (**Supplementary Fig. 2f**), using MCL-1 as a positive control as it is rapidly degraded by the proteasome. When focusing on XIAP, which was the most differentially expressed IAP in CM-challenged cells, we observed a slower decrease in XIAP protein in CM cells compared to control cells, suggesting a lower proteasomal degradation (**Supplementary Fig. 2f**). XIAP may thus be more stable in CM cells, which might explain its higher expression following CM. CM-triggered resistance to anoikis therefore involves the pro-survival IAP proteins, which can be efficiently targeted by pre-clinically validated SMAC mimetics.

IAP are commonly described to modulate NFκB pathway, while in a positive feedback loop NFκ B regulates IAP expression ,^29 30^. This test whether CM activates NFκ B, we first assessed p65 translocation from the cytoplasm to the nucleus upon CM, and showed an increase in cells displaying nuclear p65 (**Fig. 2k**). Using two luciferase-based NFκ B reporter constructs, CM was also found to increase NFκ B transcriptional activity (**Fig. 2l, m**). To test whether NFκ B activation was required for CM-driven resistance to anoikis, we expressed in MDA-MB- BSRκ ), which is a non-degradable I B that blocks the nuclear shuttling of p65 ^31^. Accordingly, the stable expression of I B^SR^ blocked p65 nuclear κ shuttling following TNFα treatment (**Supplementary Fig. 2g**). However, in these settings, blocking the NFκB pathway in CM cells did not prevent their resistance to anoikis (**Fig. 2n, o**). In addition, the artificial activation of NFκ B pathway using TNFα treatment, can upregulate cIAP2 as previously described, yet it had no effect on both cIAP1 and XIAP (**Supplementary Fig. 2h, i**) ^29^. These data suggest that CM-driven mechanical stress is characterized by NFκ B activation, which is not involved in the survival advantage observed in CM-challenged cancer cells.

To conclude, resistance to anoikis driven by CM mechanical stress relies on the pro-survival function of IAPs, regulated at the post-transcriptional level following mechanical stress.

### Confined migration enhances the aggressiveness of breast cancer cells and promotes evasion from immune surveillance

To gain mechanistic insights into the relationship between cellular constriction and resistance to anoikis, we performed an RNA sequencing (RNA seq) analysis on MDA-MB- 231 cells undergoing CM, compared to control cells. Remarkably, CM cells displayed an almost global inhibition of transcription, making their transcriptional profile distinct from control cells (**Supplementary Fig 3a, b**). To investigate this effect further, we performed a Western blot analysis for histone H3 epigenetic modifications, associated with transcriptional activation (H3K27 acetylation) or heterochromatin (H3K9 tri-methylation). Consistently with their overall transcriptional inhibition, CM cells had a lower histone H3 acetylation and a reduction in heterochromatin (lower H3K9 me3), which may indicate a decrease in nuclear stiffness, which is needed when cells navigate through narrow spaces (**Supplementary Fig. 2c**). When querying the Gene Ontology (GO) Biological Processes using both Enrichr (**Supplementary Fig. 3d**) and g:Profiler (**Supplementary Fig. 3e**), several pathways associated with cellular motility such as “Extracellular matrix organization”, “Cell-matrix adhesion” or “Cell adhesion” were significantly overrepresented in CM cells ^32 33^. Given that metastatic cells acquire an aggressive phenotype, we thus hypothesized that CM may impact on cancer cell motility ^3^.

**Figure 3:**
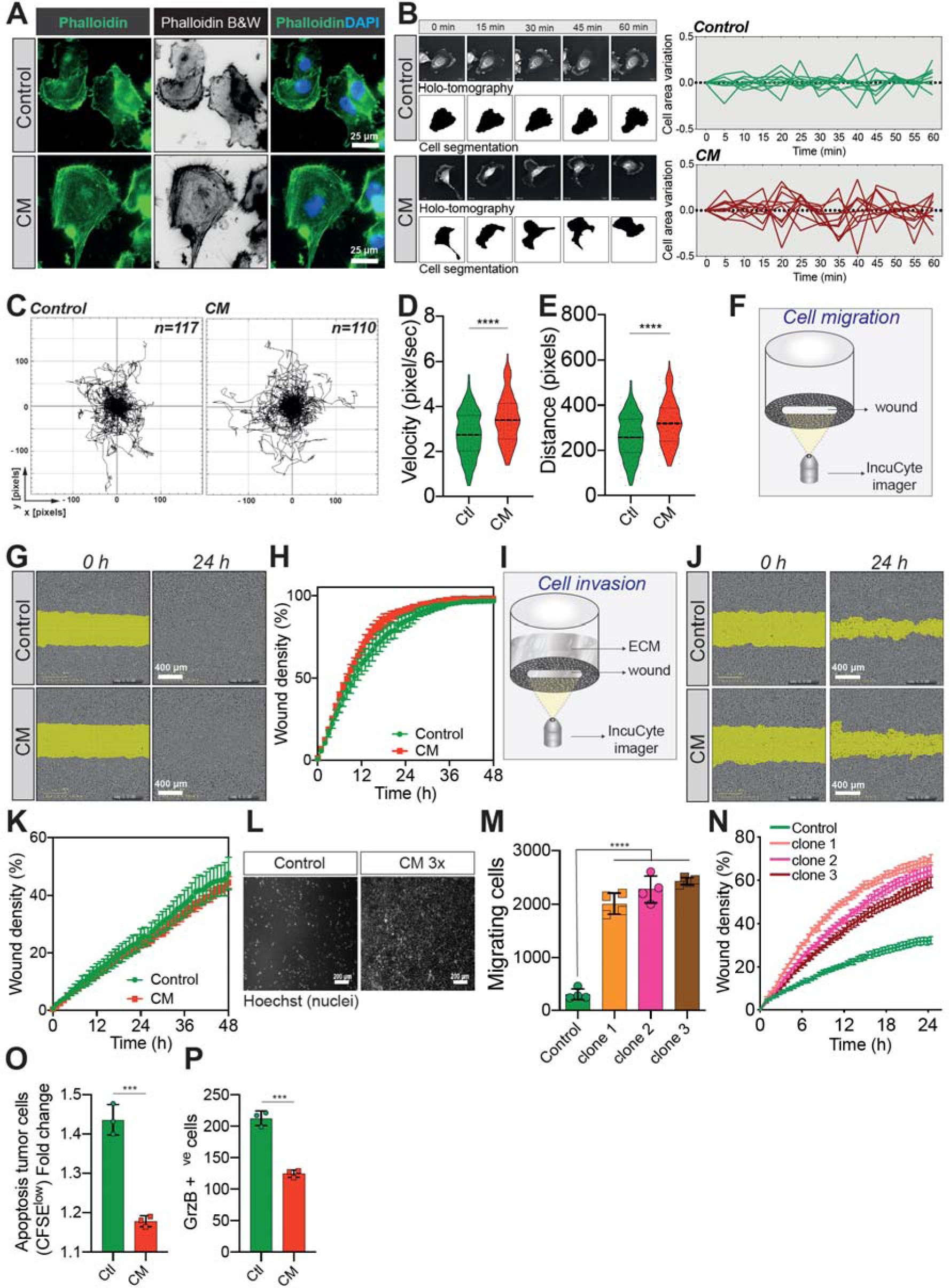
Confined migration confers breast cancer cells with a discrete aggressive behaviour. a) Representative immunofluorescence images of MDA-MB-231 cells after CM with Phalloidin-stained actin filaments. b) Representative kinetics holotomographic images (phase and cell segmentation) of control and MDA-MB-231 cells subjected to CM obtained by Nanolive imaging (left panel). Corresponding quantitative analysis of cell area variations of control and constricted cells during a 60 min time-lapse acquisition is shown in the right panel. c) Spider plot analysis for random migration assay. Control (117 cells) and CM (110 cells) cells were tracked for 24 h. d-e) Quantitative analysis of random cell migration velocity (d) and distance travelled (e) between control and CM MDA-MB-231 cells. f) Schematic representation of the IncuCyte ZOOM imager-based wound healing assay. g) Monolayers of MDA-MB-231 control and CM cells were wounded and pictures were taken immediately after wound induction (T0) and 24 h later. h) Corresponding quantitative analysis of the migratory potential of control and CM cells through wound area measurement (n=3, a representative experiment is shown). i) Schematic representation of invasion assay. After the wound was made, breast cancer MDA-MB-231 cells invaded through a matrigel plug until they closed the wound. j) Monolayers of MDA-MB-231 control and CM cells were wounded and pictures were taken immediately after wound induction (T0) and 24 h later. k) Corresponding quantitative analysis of the invasive potential of control and CM cells through wound area measurement (n=3, a representative experiment is shown). l) Representative images of control and MDA-MB-231 cells subjected to serial CM and undergoing chemotaxis through transwell membranes with 8 µm in diameter pores. m) Chemotaxis quantification relative to l) (n = 3, One-way ANOVA statistical test). n) Comparative quantitative analysis based on IncuCyte ZOOM imager of the migratory potential between control and MDA-MB-231 cells that have undergone serial CM (for three different clones) (n=3, a representative experiment is shown). o) FACS analysis of apoptotic cells among CM tumour cells compared to control cells, co- cultured with NK cells at the ratio of 1:20 for NK cells. Results were analysed by assessing the ratio of CFSE^low^/CFSE^high^ with baseline-correction to no NK cell culture condition (n=3, t test). p) FACS analysis of GrzB+ tumour cells among CM and control MDA-MB-231 cells, co- cultured with NK cells at a ratio of 1:20 (n=3, t test).

Since external mechanical forces are responsible for rapid cytoskeleton rearrangement, we first stained F-actin in control and CM-stressed MDA-MB-231 cells ^34^. This revealed a high degree of anisotropy in CM cells, with an increased number of stress fibres following CM, in addition to more abundant filopodia (**Fig. 3a**). By using Nanolive imager-based cellular tomography, we uncovered that cancer cells subjected to CM had a higher variation in cell area over time (**Fig. 3b**), consistent with variations in both cellular and

nuclear morphology, as assessed by high content screening microscopy (**Supplementary Fig. 3f**). To test whether these phenotypic changes were accompanied by increased cell motility, single-cell random migration was first tracked in control and CM stressed cells over a 24-hour period. CM-challenged cancer cells displayed a significantly higher velocity and travelled further than control cells (**Fig. 3c-e**). Proliferating cancer cells adhere to their substrate *via* focal adhesions that are equally important during migration, especially metastasis ^35, 36^. Here, we used immunofluorescence for two key components of focal adhesions, namely paxillin (**Supplementary Fig. 3g**) and vinculin (**Supplementary Fig. 3h**), to assess the number and distribution of focal adhesions following CM. The increased random migration observed was not correlated with the number of focal adhesions. In contrast to random migration, breast cancer cells subjected to a single round of CM did not outperform control cells when assessed for directional cell migration or invasion (**Fig. 3f-k**). As cells encounter several mechanical challenges during metastasis, we then imposed three consecutive CM-passages on MDA-MB-231 cells, and found that challenged cells had a significant gain in chemotaxis and directional migration (**Fig. 3l-n,** model in **Supplementary Fig. 3i**, for experimental setup).

Two of the gene expression signatures over-represented in breast cancer cells undergoing CM involve the regulation of T cell mediated immunity and most importantly the negative regulation of natural killer (NK) cells mediated cytotoxicity (**Supplementary Fig. 3d, e**). The immune system plays a crucial role in preventing metastatic dissemination through a process called immune surveillance. The innate immune NK cells and the adaptive ones, T cells αβ 37 (CD4+ and CD8+), as well as γδ T lymphocytes are the unique actors of this phenomenon ^38^. The advantage of NK-mediated immune surveillance is that it is very effective on cancer cells that are in the blood circulation ^39^. We therefore reasoned that CM might also influence NK-mediated immune surveillance. To test this, we co-cultured control and CM-challenged cells stained with CFSE with primary NK cells, obtained from healthy donor blood. Interestingly, breast cancer cells were partially protected from the NK-mediated cytotoxicity following CM (**Fig. 3o**), consistent with lower levels of granzyme B incorporation (**Fig. 3p**).

Taken together, these data demonstrate that a single event of CM has a profound effect on single-cell migration, while several rounds of CM enhance cancer cell chemotaxis and collective migration. In addition, CM contributes to evasion from NK-mediated immune surveillance.

### Breast cancer cells subjected to confined migration have an increased metastatic potential *in vivo*

We next wondered whether the effect of CM on *in vitro* breast cancer aggressiveness and invasion was applicable *in vivo*. We injected control and MDA-MB-231 cells subjected to one round of CM into the tail vein of immune-deficient mice and then analysed lung metastatic colonization by micro-computed tomography (microCT). We observed that metastasis incidence was significantly higher in the constricted cells 6 weeks post-engraftment (**Fig. 4a**). Moreover, the volume of healthy lung tissue in mice engrafted with CM breast cancer cells was considerably smaller than control counterparts, indicating their increased aggressiveness (**Fig. 4b, c**). Of note, this was also the case for breast cancer cells experiencing three consecutive rounds of CM (**Supplementary Fig. 4a**). In addition, the increased aggressiveness of constricted cells was also quantified by measuring the area of metastatic lesions following H&E staining (**Fig. 4d, e**).

**Figure 4:**
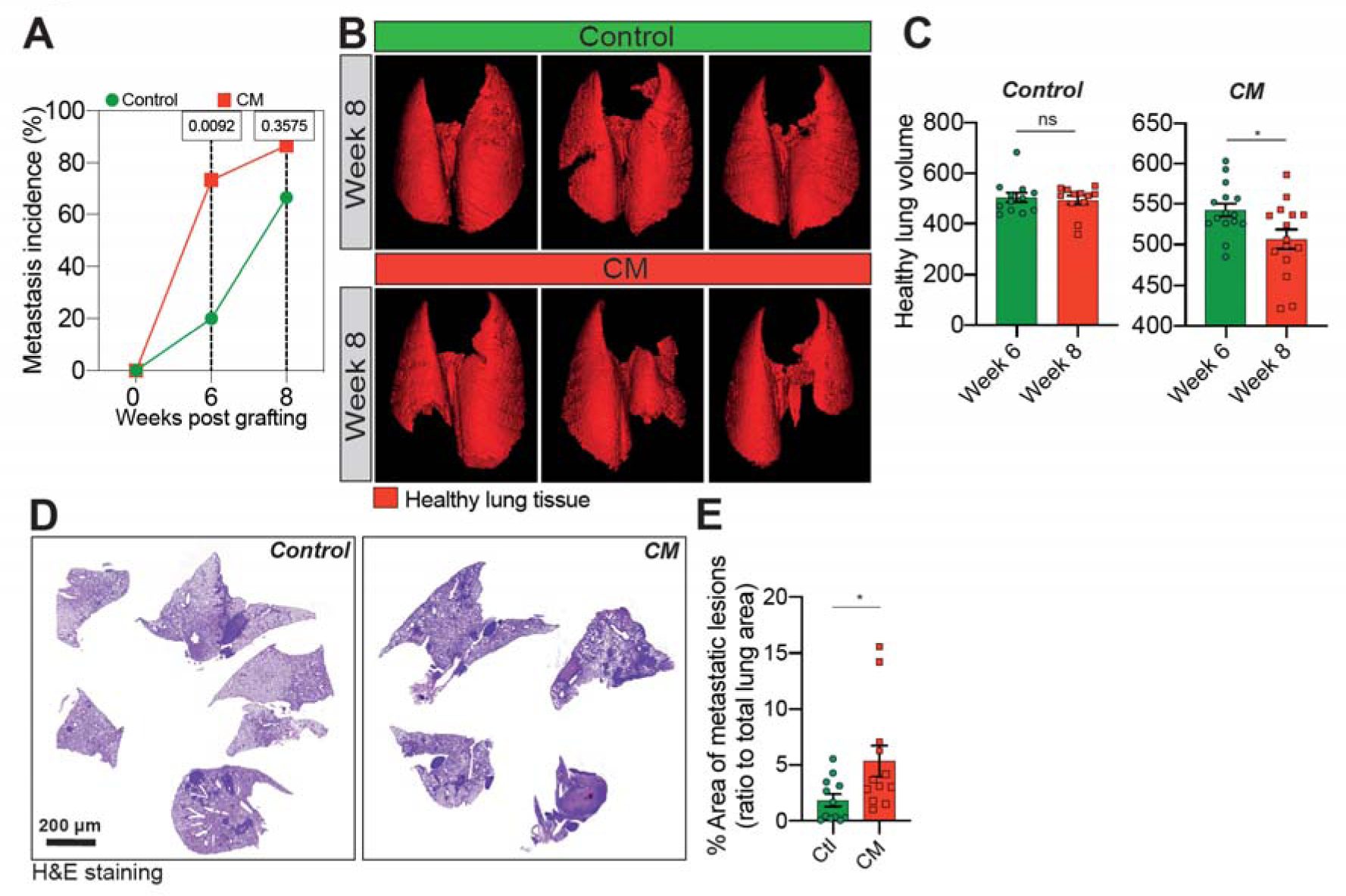
Breast cancer cells subjected to confined migration acquire an enhanced metastatic potential. a) Analysis of lung metastasis incidence in nude mice engrafted with either control or CM MDA-MB-231 cells (two-tailed Fisher’s exact test). b) Representative microCT-based 3D reconstructions (red represents healthy lung volume) of lungs from mice engrafted with either control or MDA-MB-231 cells undergoing confined migration, 15 mice/condition. c) Corresponding quantification of remaining healthy lung volume 6 and 8 weeks post- engraftment. d) Representative H&E staining of lung sections. e) Quantification of lung metastatic foci (% ratio to total lung surface).

Collectively, these data demonstrate that a single event of confined migration was sufficient to enhance lung metastatic colonization in mice. Hence, we show here that confined migration is characterised by resistance to anoikis, increased single-cell motility, activation, the resistance to anoikis relies on pro-survival IAPs, regulated at the post-transcriptional level following mechanical stress. Overall, these event contribute to enhancing breast cancer cell aggressiveness (model in **Fig. 5**).

**Figure 5:**
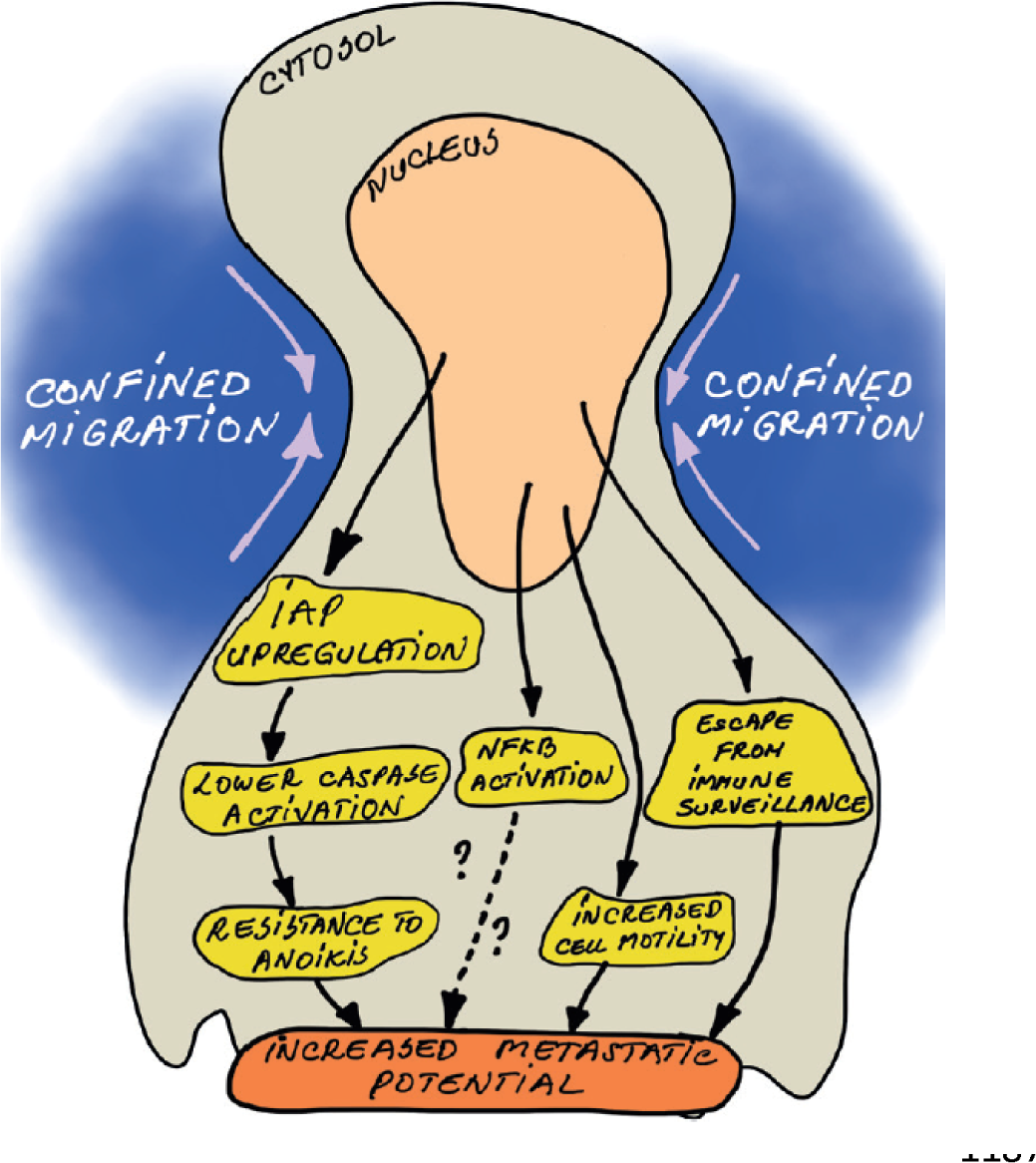
Model. As a consequence of confined migration but not compression, cancer cells become resistant to cell death triggered by loss of cell attachment (anoikis), which relies on increased expression of IAP proteins. NFkB is also activated by mechanical stress, yet it does not impact resistance to anoikis. In addition, constricted cancer cells are more resistant to Natural Killer (NK)-mediated immune surveillance. Together with a marked motility advantage, this confers an increased metastatic colonization advantage to breast cancer cells having undergone confined migration.

## Discussion

The role of cell death in cancer has been extensively investigated and its inhibition by cell-autonomous mechanisms such as overexpressing IAPs or anti-apoptotic BCL2 family proteins is a stepping stone for oncogenesis ^30 40^. However, very little is known on how the mechanically challenging tumour microenvironment impacts efficient lethal caspase activation and cancer cell death, and how this might favour cancer aggressiveness. This is a timely issue since the causal link between tumour stiffening, mechanical stress and cancer progression is well-documented ^41 42^. As the tumour stiffness and the inherent mechanical stress recently gained notoriety in favouring cancer progression, one could wonder whether this pro-oncogenic effect may be partly attributed to an underappreciated inhibitory effect on efficient induction of tumour cell death ^13 43 44^.

Here, we characterized the impact of mechanical stress, mimicking that encountered either within the primary tumour (compression) or during cancer progression (confined migration), on the acquisition of tumourigenic properties, in particular resistance to cell death. Our experimental set-up, based on commercially available transwell membranes with pores of 3 µm in diameter, forced breast cancer cells to undergo severe CM towards a chemotactic cue. Although it was previously reported that CM caused nuclear lamina breaks and widespread DNA damage, the MDA-MB-231 cells subjected to CM used herein recovered well from the forced passage, lacked obvious apoptotic caspase activation, and displayed no major differences in cell proliferation ^7 8 45 46^.

In addition to the CM that cells undergo to exit the primary tumour, CTCs also need to survive the complete loss of cell attachment, which normally triggers anoikis. Since resistance to anoikis was previously described to protect CTCs and favour their metastatic seeding, we tested this in breast cancer cells challenged by CM and unveiled a significant resistance to anoikis ^14^. As anoikis is a variation of apoptosis that relies on lethal caspase activation, we found that CM-driven resistance to anoikis was also mirrored by an inhibition of caspase activation. Although it has been reported that resistance to anoikis could be promoted by shear stress, we report here for the first time that CM, a mechanical stress in the metastatic cascade happening before the fluid shear stress, can also induce resistance to anoikis in breast cancer cells ^47^. Conversely, cancer cells migrating through larger 8 µm in diameter microporous transwells, which do not induce CM, did not acquire resistance to anoikis. Aside from constriction, cancer cells are subjected *in situ* to an important compressive stress within a rapidly growing tumour ^44^. Regarding cell death sensitivity, however, compression did not impart resistance to anoikis, thus discriminating both types of mechanical stress. Interestingly, resistance to anoikis was transient since it remained significant up to three days following the mechanical challenge. This reversibility suggests that a temporary epigenetic, transcriptional and/or translational programme is induced following acute constriction.

We then sought to uncover the cell-autonomous pro-survival pathways engaged in CM-challenged breast cancer cells and we narrowed them down to the pro-survival IAPs. Interestingly, these proteins were described to promote resistance to anoikis in several cancers. Nevertheless this is the first study showing that mechanical stress increases IAP expression, especially XIAP and cIAP1, likely by modifying their protein turnover ^24, 25^ ^26^. In an effort to revert resistance to anoikis and restore caspase-dependent cell death, we used SMAC mimetics developed to efficiently deplete IAPs. Currently, several SMAC mimetics such as birinapant and LCL161 are in phase 2 clinical trials for ovarian cancer and myeloma (source: https://clinicaltrials.gov/). Importantly we found that BV6 treatment re-sensitised cancer cells subjected to CM to anoikis, reinforcing the relevance of SMAC mimetics for clinical use. Moreover, we found that breast cancer cells experiencing CM activate the NFκB pathway, which is likely due to DNA damage ^46 48^. However, our results indicate that NFκB activation is dispensable for the acquired resistance to anoikis, yet we cannot exclude additional pro-survival effects.

To gain a clearer insight into CM-associated gene regulation, control and cancer cells experiencing CM were analysed by RNA sequencing. CM had a dramatic effect of overall transcriptional inhibition, which is most probably the immediate effect of DNA damage ^49^. This was mirrored by the marked reduction of acetylated histone H3, illustrating global transcriptional inhibition. However, the duration of inhibition following acute cell constriction remains to be investigated. In addition, it would be relevant to test by ChIP-Seq the exact gene regulatory networks impacted by loss of histone acetylation in promoter regions.

Since several gene expression signatures in CM cells were focused on cell adhesion and extracellular matrix disassembly, we next established whether breast cancer cells acquired migratory and invasive properties after recovering from a single round of CM. In line with published data, these cells displayed modifications of their cellular motility, with most obvious differences observed for F-actin filament remodelling and random migration ^11 50 51^. These data support the existing relationship between mechanical stress, the remodelling of F-actin filaments and increased motility ^35 52^. When CM was applied three consecutive times, the recovered cells were very aggressive, possibly due to the acquisition of a more stable aggressive transcriptional and/or epigenetic programme. This illustrates that multiple CM events encountered by a cancer cell during its dissemination to distant organs can have dramatic effects on its aggressive phenotype. Aside from the capacity to overcome anoikis and discrete modifications in motility, cancer cells experiencing a single CM event were also protected from NK-mediated immune surveillance. Despite these promising results, further research should be undertaken to evaluate NK function and analyse NK cytokine secretion in the presence of mechanically stressed cells. Since NK immune surveillance is conditioned by the expression on the cancer cell surface of MHC/HLA class I molecules and activation ligands, their expression should also be profiled following a mechanical stress ^39^.

Finally, we tested whether CM-triggered resistance to anoikis, increased single cell motility and evasion from NK-mediated immune surveillance were reflected *in vivo* by increasing the metastatic potential of invading cells. This was indeed the case since MDA-MB-231 breast cancer cells subjected to a single round of CM had a significant advantage to form metastatic lesions in the lungs of nude mice. This was also evidenced for cancer cells experiencing several rounds of CM.

In summary, this study refines our understanding of the pathophysiological relationship between mechanical stress and cancer aggressiveness. Our model of mechanical stress mimicking confined migration had an unexpected effect on resistance to anoikis, which was then mirrored by an enhanced metastatic seeding. In addition, our findings unveiled a previously unknown reliance of mechanically-challenged breast cancer cells on IAPs for survival that could be targeted by treatment with SMAC mimetics.

## Material and methods

### Cell lines

The cancer cell lines were obtained from American Type Culture Collection (ATCC). Human breast cancer cells MDA-MB-231 and Hs 578T were maintained in RPMI supplemented with 2 mM L-glutamine (ThermoFisher Scientific, 25030-24), non-essential amino acids (ThermoFisher Scientific, 11140-035), 1 mM sodium pyruvate (ThermoFisher Scientific, 11360-039), 10% FBS (Eurobio, CVFSVF00-01) and 1% penicillin/streptomycin (ThermoFisher Scientific, 15140-122).

### Stable cell line generation by lentiviral transduction

293T cells (1.5 x 10^6^ in a 10 cm Petri dish) were transfected with lentiviral plasmids together with pVSVg (Addgene, 8454) and psPAX2 (Addgene, 12260) using Lipofectamine 2000 (ThermoFisher Scientific, 11668019) according to the manufacturer’s instructions. Twenty-four and 48 h later, virus-containing supernatant was collected, filtered, supplemented with 1 µg/mL polybrene (Sigma-Aldrich, H9268) and used to infect target cells. Two days later, the transduced cells were selected by growth in the appropriate antibiotic.

### Plasmid transfection

For the transient overexpression of cIAP1 and XIAP, MDA-MB-231 cells (1.2 x 10^6^) were plated overnight on a 10 cm petri dish. The cells were then transfected using Lipofectamine 2000 (ThermoFisher Scientific, 11668019) with pcDNA3 as empty vector or PEF-hXIAP-Flag for XIAP. A co-transfection with PV2L-Blasti-TRAF2 and pEF6 2xHA cIAP1 WT was needed for overexpressing cIAP1. Six hours later, the transfection medium was replaced by fresh medium and the cells were allowed to grow for 48 h.

### Generation of CRISPR/Cas9-based KO cells

The oligos containing the gene-specific sgRNA target were cloned into the LentiCRISPRv2 Blasticidin (Addgene, 83480) as previously described ^53^. Following lentiviral transduction, cells were selected with 10 mg/mL blasticidin (Invivogen, ant-bl) for 2 weeks prior to analysis. The CRISPR/Cas9 primers are presented in the table 1 below.

**Table 1:**
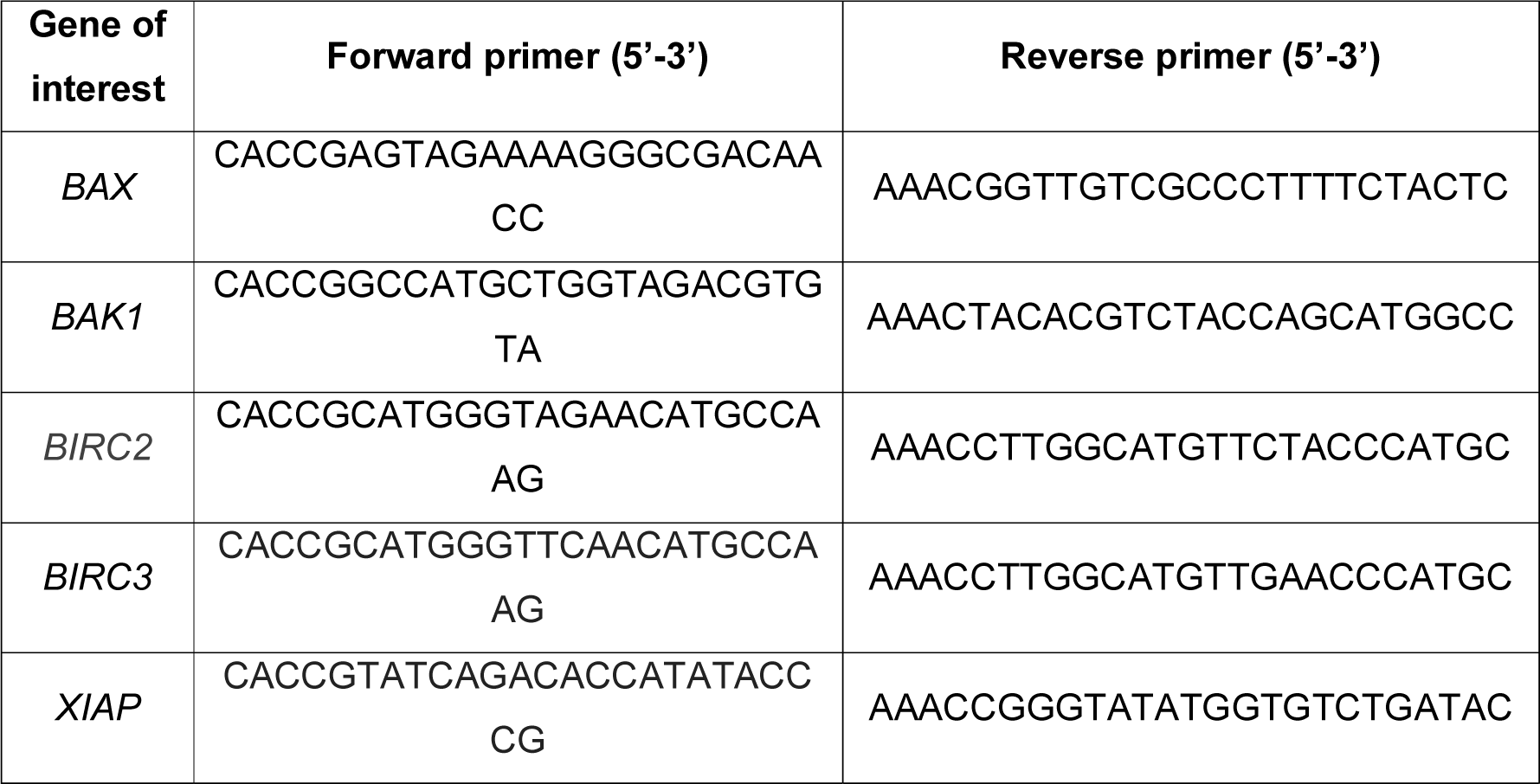
List of CRISPR primers

### Transwell assays

Breast cancer cells (MDA-MB-231, Hs578 t; 3 x 10^6^) were plated on a 75 mm transwell insert with a polycarbonate membrane pore size of 3 µm (Corning, 3420). Before seeding, the insert was coated with a layer of matrigel (300 µg/mL). A gradient of serum was then created between the two compartments of the transwell (0% FBS in the top compartment and 20% below) and renewed 5 h later. After 72 h, cells were harvested after washing the insert with PBS and incubation with trypsin. Cells that have migrated through the 3 µm pores were designated as the “constricted cells”, whereas the ones that did not migrate were the “control cells”.

For the 8 µm transwell experiments, 5 x 10^5^ MDA-MB-231 cells were plated on a 6-well insert with a polycarbonate membrane pore size of 8 µm (Greiner Bio One 657628) and the cells were recovered and analysed 48 h later.

For certain experiments, cells recovered for the transwell assay were stained with Hoechst 33342 (10 µg/mL, ThermoFisher Scientific, H1399), calcein AM (0.4 mg/mL, Life, C1430) and Vybrant ™ cell-labelling solutions (DiI and DiO, V-22885 and V-22886) according to the manufacturer’s instruction.

### Anoikis assay

MDA-MB-231 and Hs578T cells (2 x 10^5^) were seeded onto a 6-well Clear Flat Bottom Ultra-Low Attachment plate (Corning, 3471) in complete RPMI medium. Cells then formed 3D clonogenic structures that were imaged and scored after 7 and 14 days of culture. Six wells were plated for each condition and experiments were repeated three times for each cell line. Cells were also grown on a 1 % agarose petri dish in RPMI medium without serum for 24 h and proteins were extracted with protein lysis buffer.

### Soft agar colony assay

Cells (10^3^/well) were suspended in 1 mL of 0.3% low gelling temperature agarose (Sigma, A9414) and plated onto a 1% agarose layer in three wells of a 6-well plate. When the 0.3% agarose solidified, the wells were covered in complete RPMI media and colonies were scored 4 weeks later.

### Operetta CLS High-Content Analysis

Control and constricted MDA-MB-231 cells (1.5 x 10^4^) were seeded onto 8 wells of a 96-well plate. Twenty-four hours later, the cells were stained with 0.4 mg/mL calcein AM and 2 µg/mL Hoescht (ThermoFisher Scientific, H1399), for 30 min at 37°C. The medium was then replaced with phenol red-free DMEM (ThermoFisher Scientific, 21063029) and the cells were imaged with the Operetta CLS (PerkinElmer), and the cell morphological analysis was performed using the Columbus software (PerkinElmer).

### Immunofluorescence

MDA-MB-231 (5 x 10^4^) were seeded onto coverslips placed in 24-well plate overnight. For studies on focal adhesions, the coverslips were first coated with 100 µg/mL matrigel (Sigma Aldrich, E1270). After washing in PBS, cells were fixed in 4% PFA for 5 min and then washed once. Cells were permeabilized with 0.2% Triton X-100 (Pan Reac, A4975.01) diluted in PBS, for 10 min at room temperature and the blocking of non-specific binding sites was done using 2% BSA in PBS, for 1 h at room temperature. Cells were then incubated with the primary antibody COX IV (Cell Signaling, 4850S), cytochrome *c* (Cell Signaling, 12963S), Alexa Fluor 647 Phalloidin (Invitrogen, A22287), vinculin (Sigma-Aldrich, V9131), paxillin (BD Transduction Biosciences, 610052) or p65 at 1/400-500 dilution in PBS, for 1 h at room temperature or overnight at 4°C. Next, the cells were washed in PBS three times and then incubated with the appropriate secondary antibody coupled to Alexa Fluor (1/300, Thermofisher scientific, A21151 and A31571) for 1 hour at room temperature protected from light. The staining of nuclei was done with Hoechst 33342 (10 µg/mL, ThermoFisher Scientific, H1399) or with DAPI mounting medium (Vectashield). The coverslips were finally mounted using Fluoromount (Southern Biotech, 0100-01). Slides were left to dry overnight before image acquisition using a Zeiss Axio Imager microscope (Zeiss).

### Anoikis resistance assay using the IncuCyte ZOOM imager

MDA-MB-231 cells (10^4^ cells) were plated in a 96-well Clear Round Bottom Ultra-Low Attachment Microplate (Corning, 7007). SytoxGreen (30 nM, Life, 1846592) was also added to the medium to stain apoptotic cells. Cells were then imaged every 60 min using the IncuCyte ZOOM imager.

### VC3AI reporter-based caspase activation assay

MDA-MB-231 VC3AI (control and constricted) cells (2 x 10^5^) were collected from transwell assay and the mean fluorescence intensity of the green signal (VC3AI) was then determined by FACS Calibur flow cytometry (BD Biosciences, San Jose, CA, USA). Control cells were treated with 1 µM Actinomycin D as a positive control for cell death.

### Fluorometric caspase 3/7 activity assay

The cell pellets were resuspended in Cell Lysis Buffer before evaluating the amount of protein in each sample. Twenty µg of proteins were then mixed with Reaction Buffer supplemented with DEVD-AFC substrate. After 1 h of incubation at 37°C, the caspase 3/7 activity was assessed by fluorescence measurement. Caspase 3 activity was determined using the Caspase 3/CPP32 Fluorometric assay kit according to the manufacturer’s instructions (BioVision, K105).

### Mitochondrial membrane potential assay

MDA-MB-231 cells (10^5^) were harvested and resuspended in 0.1 µM Tetramethylrhodamine Ethyl Ester Perchlorate (TMRE, ThermoFischer Scientific, T669) for 30 min at 37°C. This fluorescent compound accumulates only in intact mitochondria and highlights the mitochondrial membrane potential of living cells. CCCP (carbonyl cyanide 3- chlorophenylhydrazone) was used as a mitochondrial membrane potential disruptor. When mitochondria are depolarized or inactivated, leading to a decrease in membrane potential, TMRE accumulation is reduced. After washing, the membrane potential (mean fluorescence intensity of the red signal) was determined by flow cytometry.

### Evaluation of mitochondrial superoxide levels

MDA-MB-231 VC3AI (control and constricted) cells (5 x 10^4^) were harvested and resuspended in 5 µM Mitosox (ThermoFischer Scientific, M36008) for 15 min at 37°C. After washing with PBS, the level of mitochondrial superoxide was determined by flow cytometry.

Cells treated 1 h with 500 µM H_2_O_2_ were used as a control for the production of reactive oxygen species (ROS).

### Measurement of total ROS in live cells using CellROX staining

After plating 5 x 10^4^ MDA-MB-231 cells (control and constricted) overnight in a 12-well plate, cells were trypsinized and treated 30 minutes at 37°C with 5 µM CellROX Deep Red reagent (Life technologies, C10422) diluted in medium. This cell-permeant dye exhibits a strong fluorescence once oxidized by cytosolic ROS. Positive control of ROS consisted in cells treated 1 h with 500 µM H_2_O_2_ (Sigma, H1009) before staining. Cells were then centrifuged and washed in PBS three times, before being resuspended in 200 µL of medium. The subsequent analysis was performed using FACS Calibur.

### ATP assay

ATP levels were measured in control and constricted breast cancer cells (3 x 10^5^ cells) using the ATP fluorometric assay kit (Sigma-Aldrich, MAK190), following the manufacturer’s instructions. Cells treated 1 hour with 500 µM H_2_O_2_ served as a negative control. To evaluate the ATP concentration, an ATP standard curve from 0 to 10 nM was used.

### Cell cycle analysis

10^5^ MDA-MB-231 cells (control and constricted) were washed once in PBS and pelleted in FACS tubes. Cold ethanol at 100% was added drop by drop while vortexing at a final concentration of 70% in PBS. Cells were then stored at -20°C until use. For FACS analysis, the cells were rinsed with PBS then centrifuged at 850 g for 5 min. The pellet was then treated with 100 µg/mL ribonuclease A (A8950) in order to specifically stain DNA. Propidium iodide (Sigma, P4864) was added at 100 µg/mL and cells were immediately analysed by flow cytometry.

### Cell compression

MDA-MB-231 cells (3 x 10^6^) were plated on 75 mm transwell inserts with a polycarbonate membrane pore size of 3 µm (Corning, 3420) that allows media and gas exchange during compression. Twenty-four hours later, a 2 % agarose (Sigma, A9539-100G) disk was placed on top of the cells in order to prevent the direct contact with the plastic cup (3D printed by F. B.) placed above the agarose disk. A range of pressure was then tested for 24 h (0, 200, 300, 400, 600 Pa) by adding the appropriate lead weights in the plastic cup. At the end of the compression time (24 h), the cells under the agarose disk were washed and collected for further analysis.

### Western blot analysis

Proteins were isolated by lysing cell pellets in RIPA lysis buffer (Cell Signaling, 9806S) supplemented with phosphatase inhibitors complex 2 and 3 (Sigma Aldrich, P5726-1ML, P6044-1ML), DTT 10 mM and protease inhibitor cocktail (Sigma-Aldrich, 4693116001). The protein concentration was then determined using the Protein Assay dye Reagent Concentrate (Biorad, 50000006). Equal amounts (15-20 μg) of each sample were separated on 4-12 % SDS-polyacrylamide gels (Biorad) under denaturating conditions (SDS PAGE Sample loading buffer (VWR, GENO786-701) supplemented with 1 mM DTT). The gels were then transferred onto a nitrocellulose membrane using the Transblot Turbo Transfer System (Biorad, 1704150EDU). An incubation of 1 h with Intercept blocking buffer (Licor, 927-70001) blocked non-specific binding sites before incubating the membranes with the primary antibody (1/1000 in Intercept T20 Antibody Diluent (Licor) overnight at 4°C, under agitation. The primary antibodies used were: actin (Sigma-Aldrich, A3854), MCL-1 (Cell Signaling, 4572S), PARP-1 (Cell Signaling, 9532), Caspase-3 (Cell Signaling, 9662S), GFP (Life, A11122), BAX (Cell Signaling, 2772S), BAK (Cell Signaling, 12105S), HSP60 (Cell Signaling, 4870), K48-Ub (Cell Signaling, 8081S), COX IV (Cell Signaling, 4850S), cIAP1 (Cell Signaling, 7065T), cIAP2 (Cell Signaling, 3130T), XIAP (Cell Signaling, 14334S), HSC70 (Santa Cruz Biotechnology, sc-7298), H3K27ac (Diagenode, C15210016), H3K9me3 (Diagenode, C15200153). The membranes were rinsed 4 times for 5 min in TBST 0.1% and then incubated with appropriate secondary antibody coupled to IRDye® 800CW or 680RD dye (Licor; 1/10000) for 1 h at room temperature under agitation and protected from light. Four extra washing steps in TBST 1% and 1 in TBS were performed before scanning the membrane by Odyssey® Imaging System for near infrared detection.

### Cycloheximide chase assay

To determine protein half-life, control and constricted MDA-MB-231 cells (5 x 10^5^) were treated in ultra-low attachment conditions with cycloheximide (CHX; 50 µg/mL) for different durations (0, 6, 16, 24, 33, 48 h) and protein extracts were analysed by Western blot.

### Dual luciferase reporter assay

MDA-MB-231 cells (10^5^) were plated in 12-well plates for 24 h and were then co-transfected with the NFkB luciferase reporter-containing plasmid and a Renilla plasmid using Lipofectamine 2000. After 48 hours of transfection, the luciferase activity was assessed with the Dual luciferase reporter assay (Promega, E1910) following the manufacturer’s instructions. Firefly luciferase activity was then normalized against Renilla luciferase activity.

### Holotomographic microscopy (HTM)

MDA-MB-231 cells (5 x 10^4^) cells were seeded onto Fluorodishes (ibidi GmbH, Gräfeling, Germany). HTM was performed on the 3D Cell-Explorer Fluo (Nanolive, Ecublens, Switzerland) using a 60X air objective at a wavelength of λ = 520 nm. Physiological conditions for live cell imaging were maintained using a top-stage incubator (Oko-lab, Pozzuoli, Italy). A constant temperature of 37°C and an air humidity saturation as well as a level of 5% CO_2_ were maintained throughout imaging. Refractory index maps were generated every 5 min for 1 h. Images were processed with the software STEVE.

### Random migration assay

10^3^ breast cancer cells were seeded onto a 96-well ImageLock plate (Sartorius, 4379) and imaged for 24 hours using the IncuCyte ZOOM-based time-lapse microscopy. The acquired time-lapse images were treated with a manual tracking plugin using the Image J software. About 100 cells/condition were followed for 30 min in order to determine the accumulated distance and their velocity.

### Wound healing assay

MDA-MB 231 cells (5.5 x 10^4^) were seeded onto a 96-well imageLock plate (Sartorius, 4379) and grown for 24 h until cell confluency was reached. A scratch was then performed in the cell monolayer using a WoundMaker (Sartorius, 4563), following the manufacturer’s instructions. Wound closure was imaged and quantified using the IncuCyte ZOOM imaging system.

### Invasion assay

Wells of a 96-well imageLock plate (4379, Sartorius) were first coated with 100 µg/mL of Matrigel (Sigma Aldrich, E609-10mL). After 1 h, MDA-MB-231 cells (5.5 x 10^4^) were seeded 24 h prior to the assay. A wound was then performed in the cell monolayer with the WoundMaker and a new layer of Matrigel (800 µg/mL) was deposited onto cells for 1 h at 37°C to allow polymerization. The top of the cells was covered with complete medium and the invasion potential of cancer cells was evaluated and quantified using IncuCyte ZOOM-based time-lapse microscopy.

### RNA sequencing

RNA sequencing from control and mechanically-challenged MDA-MB-231 cells was done by the CRCL Cancer Genomics core facility. The libraries were prepared from 600 ng total RNA using the TruSeq Stranded mRNA kit (Illumina) following the manufacturer’s instructions. The different steps include the PolyA mRNA capture with oligo dT beads, cDNA double strand synthesis, adaptors ligation, library amplification and sequencing. Sequencing was carried out with the NextSeq500 Illumina sequencer in 75 bp paired-end.

### Bioinformatics analysis

All genomic data were analysed with R/Bioconductor packages, R version 4.0.3 (2020-10-10) [https://cran.r-project.org/; http://www.bioconductor.org/] on a linux platform (x86_64-pc-linux-gnu [64-bit]).

Illumina sequencing was performed on RNA extracted from triplicates of each condition. Standard Illumina bioinformatics analyses were used to generate fastq files, followed by quality assessment [MultiQC v1.7 https://multiqc.info/], trimming and demultiplexing. ’Rsubread’ v2.4.3 was used for mapping to the hg38 genome and creating a matrix of RNA-Seq counts. Rsamtools v2.6.0 * was used to merge two bam files for each sample (run in two different lanes). Next, a DGElist object was created with the ’edgeR’ package v3.32.1 [https://doi.org/10.1093/bioinformatics/btp616]. After normalization for composition bias, genewise exact tests were computed for differences in the means between groups, and differentially expressed genes (DEGs) were extracted based on an FDR-adjusted p value < 0.05 and a minimum absolute fold change of 2. All raw and processed RNAseq data have been deposited at the Gene Expression Omnibus (GEO) repository, under accession number GSE176081.

* Martin Morgan, Hervé Pagès, Valerie Obenchain and Nathaniel Hayden (2020). Rsamtools: Binary alignment (BAM), FASTA, variant call (BCF), and tabix file import. R package version 2.6.0. https://bioconductor.org/packages/Rsamtools

### Quantitative RT-PCR

Total RNA extraction was performed using the Nucleospin RNA Macherey-Nagel kit (740955) and quantified by NanoDrop. The conversion of messenger RNA into cDNA was performed using the Sensifast cDNA synthesis kit (Bioline, BIO- 65053). cDNA was then amplified by PCR using specific primers for each gene designed with Primer-blast software (https://www.ncbi.nlm.nih.gov/tools/primer-blast/) and listed in Table 2. GAPDH, ACTB and HPRT were used as housekeeping genes. The thermal cycling steps included an initial polymerase activation step at 95°C for 2 min, followed by 40 cycles at 95°C, 5 s, and 60°C, 30 s. The qRT-PCR experiments were performed using SYBR Green and a Lightcycler96 (Roche, Indianapolis, USA).

**Table 2:**
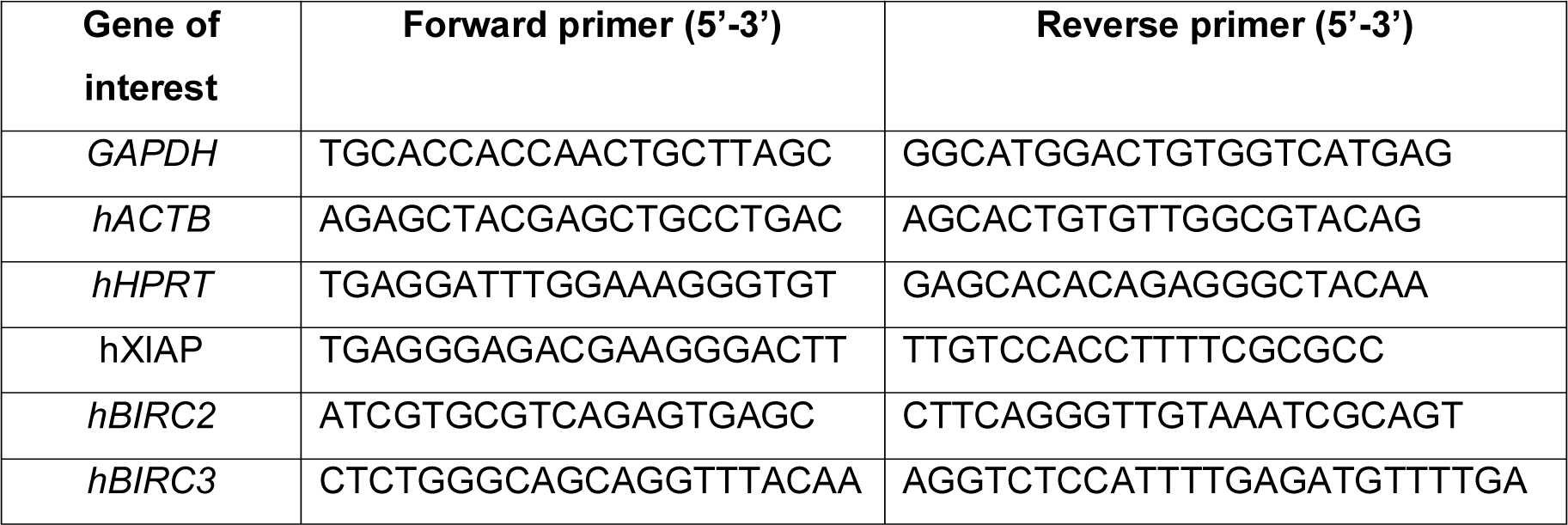
List of qRT-PCR primers

### In vivo lung metastasis model and lung imaging

MDA-MB-231 control and constricted cells (2.5 x 10^4^) were suspended in 100 µL PBS and injected into the tail vein of NMRI nude female mice. The absence of mycoplasma in injected cells was controlled before injection in animals. Burden of lung metastasis was evaluated over time by X-ray microCT-Scan (Quantum FX®, Perkin Elmer). Mice were anesthetized with a continuous flow of 2 to 4% isoflurane/air (1.5 L/min). The lungs were imaged in a longitudinal manner for 2 min with an exposure of 0.746 Gy and the obtained raw data were reconstructed with the following acquisition settings: a 24mm FOV diameter, 512 slices and 50 µm voxel. The resulting images were viewed and analysed using “Analyze of Caliper” software (AnalyzeDirect) and the remaining healthy lung volume was quantified and 3D- represented.

### Histological analyses

If limiting points were not observed, mice were euthanized 8 weeks post-engraftment. Lungs were fixed in 4% buffered formalin, paraffin embedded and three 3 µm-sections separated by 300 µm were stained with Hematoxylin-Eosin. The slides were scanned using the panoramic scan II (3D Histech). These were then analyzed with CaseViewer 2.2.0.85100 software (3DHISTECH Ltd.) for the detection of metastasis.

### NK-mediated immune-surveillance

Control and constricted MDA-MB-231 cells were cocultured with human NK cells sorted from peripheral blood using the NK cell isolation Kit (Miltenyi Biotec 130-092-657) at the ratio of 1:20, in triplicate for each condition. Before co-culturing, tumour cells were pretreated with 10 µg/mL mitomycin for 1 h to stop proliferation, and were incubated and tagged with CFSE (Invitrogen CellTrace, C34570) at 1 µL/mL for 20 min. Twenty-four hours later, cells in each condition were recovered by trypsin and stained intracellularly with Granzyme B (Biolegend, AF647, clone GB11). Flow cytometry was performed by BD LSR Fortessa HTS and data were analysed using GraphPad Prism V9.

### Image analysis

Image analysis was performed using the ImageJ software 1.52a.

### Statistical analysis

Data are expressed as the mean ± SEM. A two-tailed Student’s t-test was applied to compare two groups of data. Analyses were performed using the Prism 5.0 software (GraphPad).

## Acknowledgements

This work was supported by funding from LabEx DEVweCAN (University of Lyon), Agence Nationale de la Recherche (ANR) Young Researchers Project (ANR-18-CE13-0005-01), La Ligue Nationale Contre le Cancer and Fondation de France. We thank Brigitte Manship for reviewing the manuscript, Virgile Raufaste-Cazavieille, Léa Magadoux and Thomas Barre (AniCan Image, Lyon, France, funding PHENOCAN ANR -11-EQPX-0035 PHENOCAN) and the Anatomopathology core facility for technical assistance.

## Author contributions

<u>Conceptualization</u>, G. Ichim, S.T. and D.F.; <u>Methodology</u>, G. Ichim, D.F., H. H.-V.; <u>Formal</u> <u>analysis</u>, G. Ichim, D.F., Z. W., H. H.-V. and A. H.; <u>Investigation</u>, G. Ichim, D.F., Z. W., J. M., K. B., D. N.,K. W., A. H. and H. H.-V.; <u>Resources</u>, D. N., F. B. and A. H.; <u>Writing</u> – Original Draft and Editing, G. Ichim and D. F.; All authors reviewed and edited the manuscript; <u>Supervision</u>, G. Ichim, D. F. and A. H.; <u>Project administration and funding acquisition</u>, G. Ichim.

---

**Supplementary figure 1.**
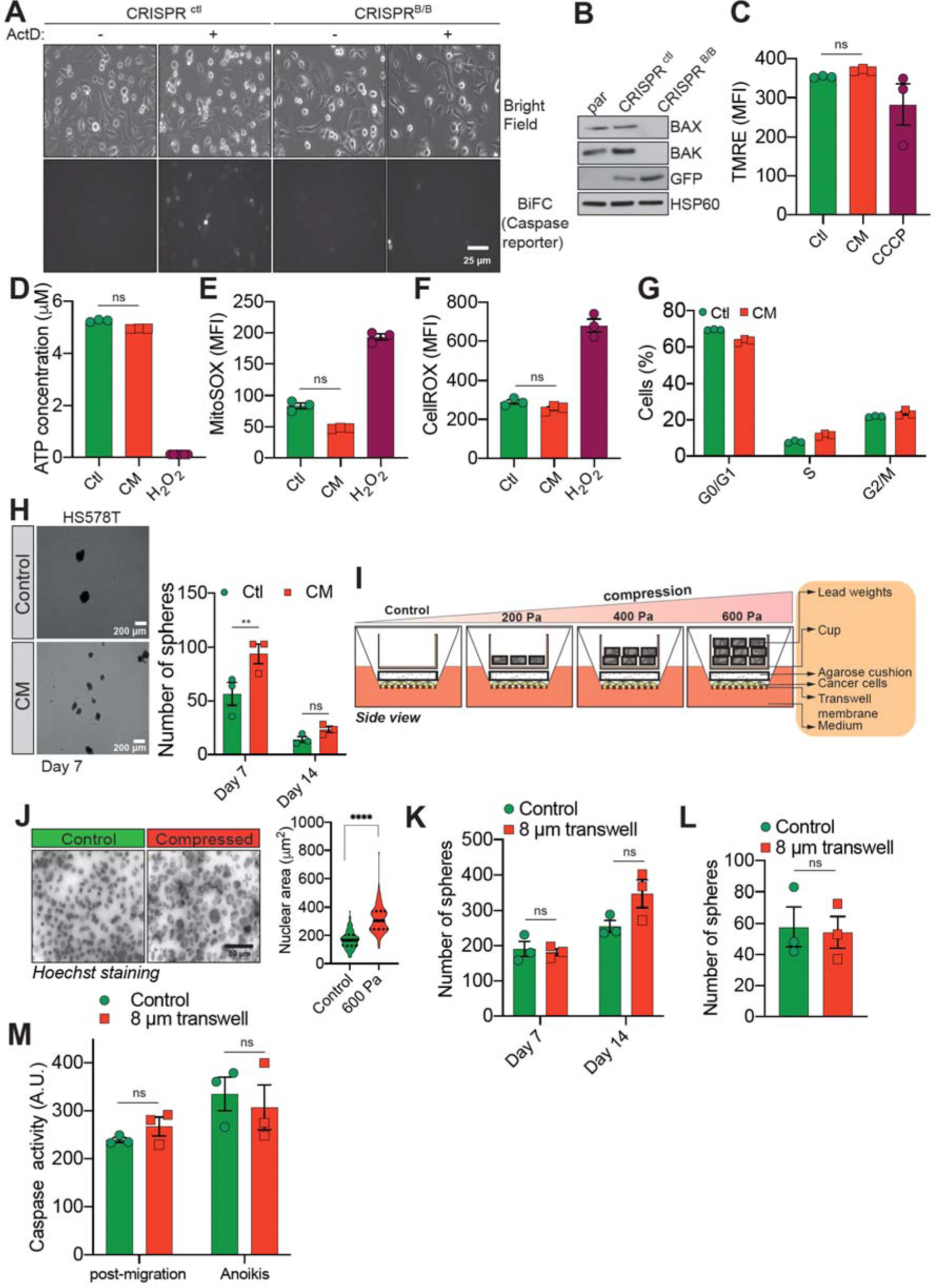
(related to Figure 1) a) Representative epifluorescence and phase images of VC3AI caspase reporter-expressing control and BAX/BAK double-KO MDA-MB-231 cells validating the BiFC caspase reporter. Treatment with Actinomycin D was used as a positive control to induce caspase activation. b) Western blot analysis of BAX and BAK protein expression, validating their knocking out through CRISPR/Cas9 in VC3AI-expressing (GFP positive) MDA-MB-231 cells. c) Flow cytometric analysis of mitochondrial membrane potential (TMRE staining). CCCP treatment at 12.5 µM was used as a positive control for disrupting the mitochondrial membrane potential (n = 3, One-way ANOVA statistical test). d) Quantification of total ATP content in control and constricted MDA-MB-231 cells (n = 3, One-way ANOVA statistical test). e-f) Flow cytometry-based quantification as mean fluorescence intensity (MFI) for mitochondrial reactive oxygen species (ROS) (e) and overall cellular ROS (f). Cells treated with 500 µM H_2_O_2_ were used as a control for the production of ROS (n = 3, One-way ANOVA statistical test). g) Assessment of MDA-MB-231 cell cycle profile using PI staining and flow cytometry analysis. h) Representative pictures of control and CM Hs578T cells cultured in ultra-low attachment condition and forming clonogenic structures (left panel). The corresponding quantitative analysis of the number of clonogenic structures after 7 and 14 days of culture is shown in the right panel (n = 3, Two-way ANOVA statistical test). i) Schematic diagram of the compression device, allowing the compression of breast cancer cells (MDA-MB-231) with a constant force. Cells are plated on a transwell membrane, allowing gas and passage of nutrient. An agarose cushion is placed on top of the cell monolayer (control cells were compressed only by the agarose disk) and a custom printed cup is then added on top and filled with the corresponding weight in lead, resulting in various pressures. j) Representative images of MDA-MB-231 nuclei (stained with Hoechst) during 600 Pa compression (left panel) and the corresponding quantification of nuclear area showing a significant increase under compression (right panel, n = 3, t test). k-l) Quantitative comparison of the number of clonogenic structures for MDA-MB-231 passing through transwells with 8 µm in diameter pores in ultra-low attachment conditions (k, n = 3, Two-way ANOVA statistical test) or in soft agar (l, n = 3, t test). m) The effect of MDA-MB-231 cells migrating through transwells with 8 µm in diameter pores on caspase-3/7 activation as assessed by a fluorometric assay. This was either tested immediately after the transwell assay or in cells that were subsequently grown 24 h in anoikis conditions (n = 3, Two-way ANOVA statistical test).

**Supplementary Figure 2.**
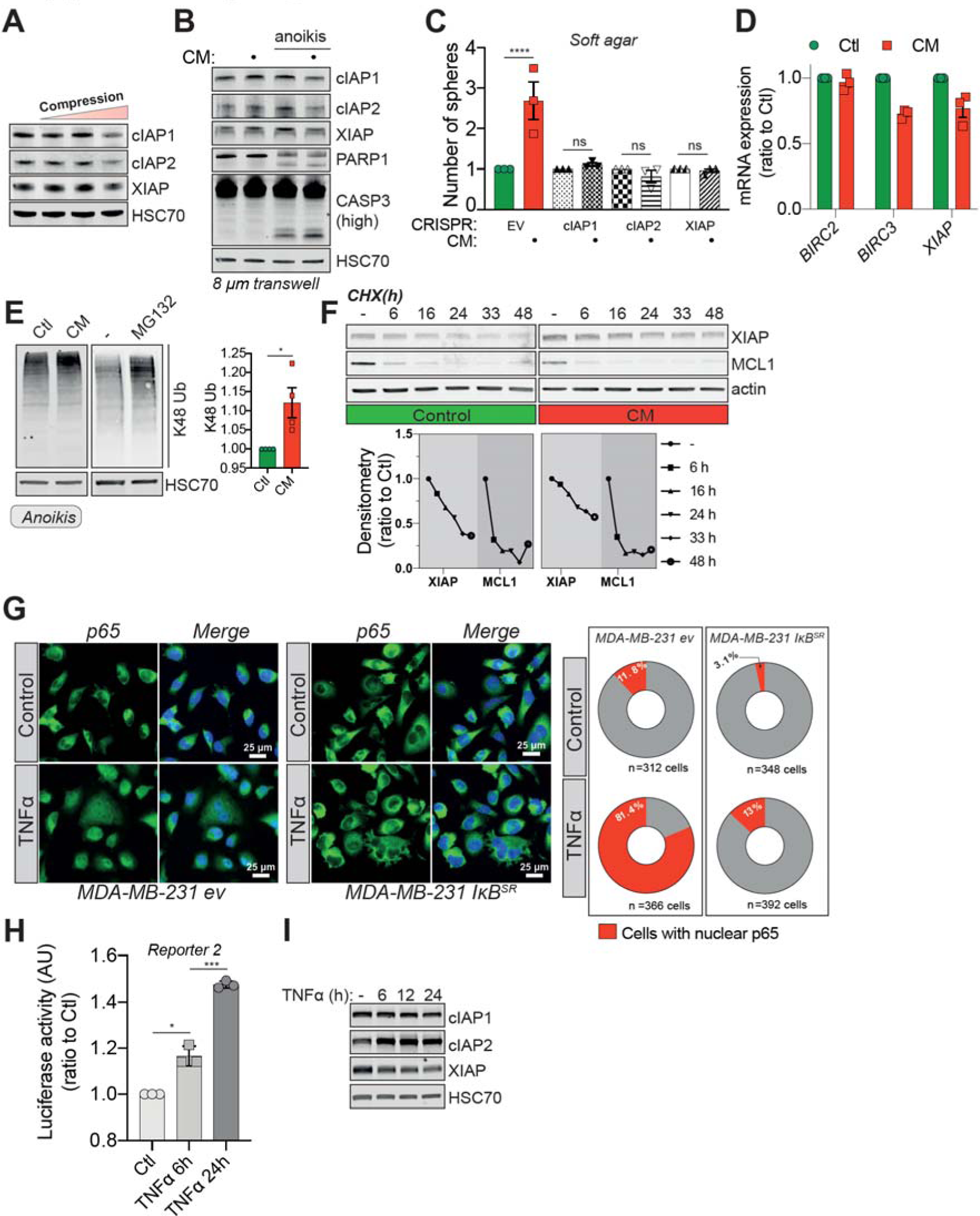
(related to Figure 2) a) Western blot analysis of IAP protein expression in MDA-MB-231 cells compressed 24 h with 200, 400 or 600 Pa. b) Western blot analysis of IAP expression, PARP-1 cleavage and caspase-3 processing in MDA-MB-231 cells after their migration through transwell pores of 8 µm in diameter. c) Quantitative comparison of soft agar clonogenic survival using control or CM MDA-MB-231 cells deleted for individual IAPs using CRISPR/Cas9 (n = 3, One-way ANOVA statistical test). d) IAP relative mRNA expression by qRT-PCR in control and CM MDA-MB-231 cells. e) Western blot analysis of K48 ubiquitination in total protein lysates from control and CM cells (left panel). Treatment with MG132 was used as a positive control for proteasome inhibition. The right panel depicts the corresponding densitometry analysis (n = 4, t test). f) Cycloheximide chase assay (50 ng/mL) showing by Western blot XIAP and MCL1 proteosomal degradation (upper panel). Lower panel represents the corresponding densitometry analysis, n = 2. g) Representative immunofluorescence images of p65 nuclear localization in MDA-MB-231 cell lines expressing the empty vector (EV) or IkB^SR^ and subjected to CM (left panel). Corresponding quantification for the percentage of cells presenting nuclear p65 staining is shown in the right panel. h) Luciferase assay validating NFkB activation following TNFα treatment (20 ng/mL) for 6 and 24 h (n = 3, One-way ANOVA statistical test). i) Western blot analysis of IAP expression in MDA-MB-231 cells treated as in N.

**Supplementary Figure 3.**
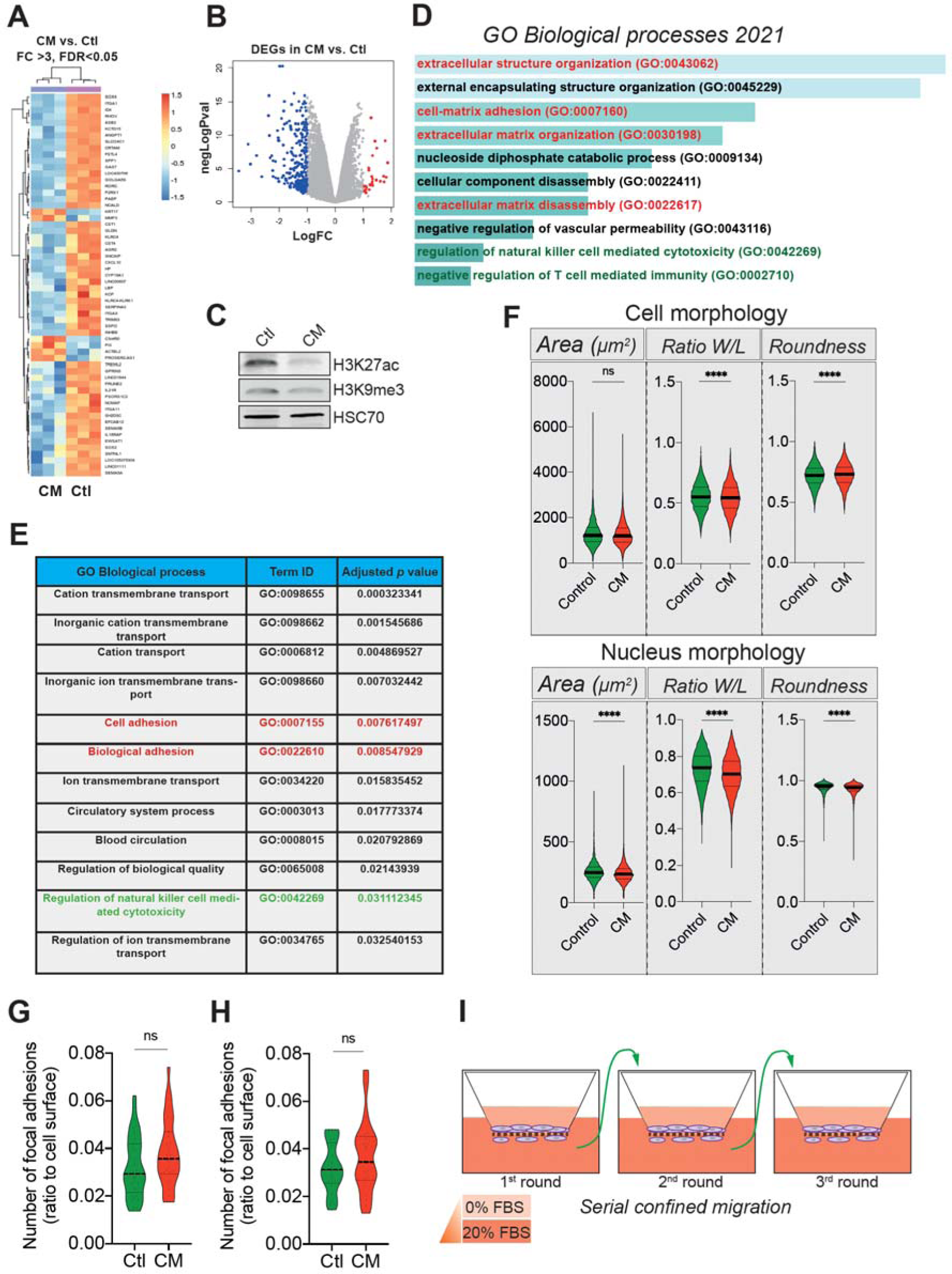
(related to Figure 3) a) Unsupervised clustering of the RNA sequencing data in control and CM MDA-MB-231 cells. Orange indicates increased and blue decreased mRNA abundance of selected genes with fold change above 3, 3 replicates/condition. b) Volcano plot displaying the expression (in log fold change) of each differentially expressed gene. c) Western blot analysis of H3K27 acetylation and H3K9 tri-methylation in control and CM MDA-MB-231 cells. d-e) Enrichr (d) and g:Profiler (e) based gene ontology (GO) analysis of the genes differentially expressed (above 2 FC, up- and downregulated) in cancer cells subjected to CM. Signatures related to cellular motility are in red while those concerning immune surveillance are in green. f) Quantitative Operetta HCS-based imaging comparison of cell and nucleus morphology parameters (area, ratio w/l and roundness). g-h) Assessment of the number of focal adhesions between control and CM MDA-MB-231 cells based on immunostaining for paxillin (g) and vinculin (h). i) Schematic representation of the serial constricted migration. Between CM events, cells were amplified and re-challenged until reaching three consecutive CM.

**Supplementary Figure 4.**
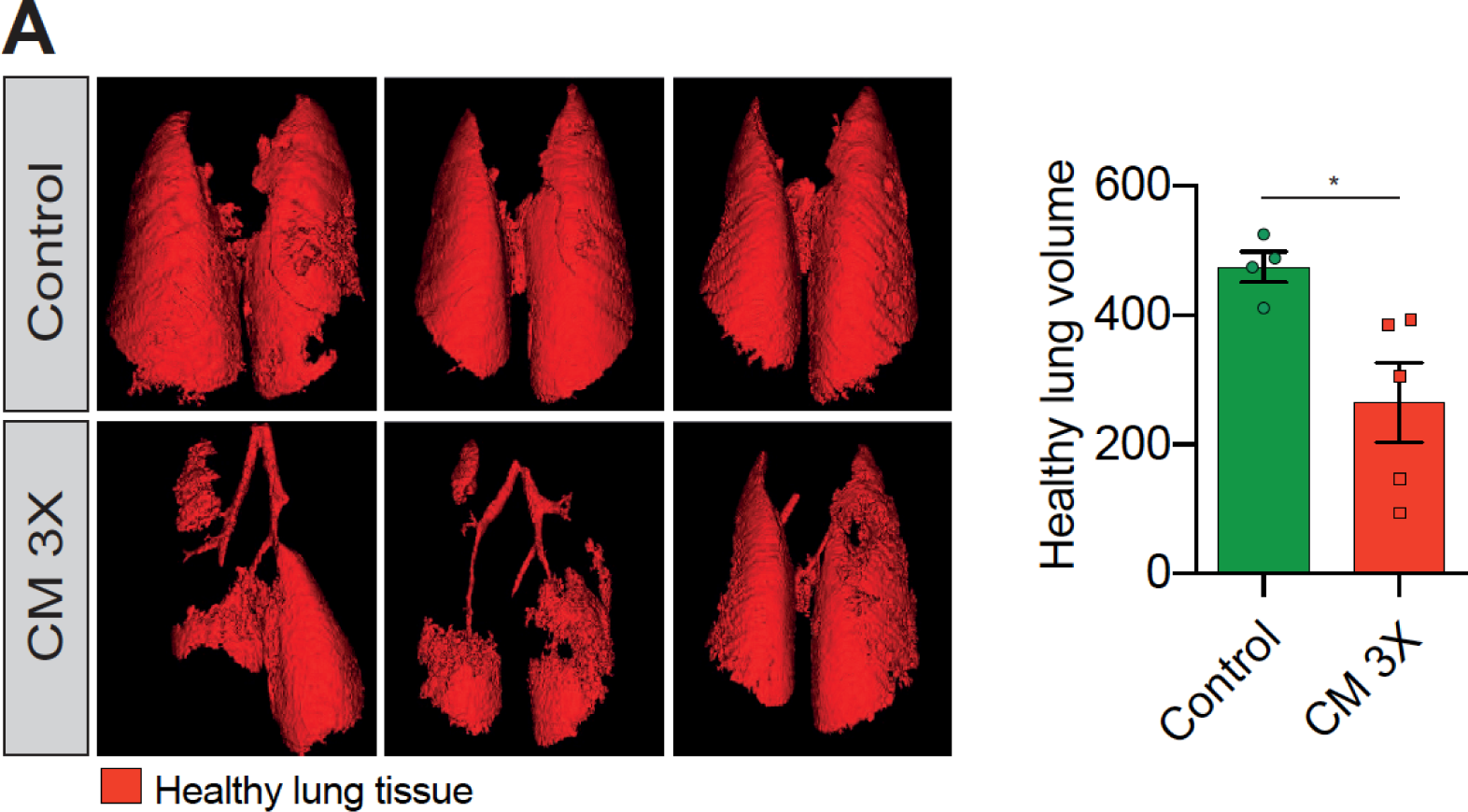
(related to Figure 4) a) Analysis of lung metastasis incidence in nude mice engrafted with either control or MDA-MB-231 cells undergoing three consecutive rounds of CM (left panel). Right panel presents the corresponding quantification of remaining healthy lung volume in engrafted mice, 4-5 mice/condition.

